# Recombination and the Species Structure of the Genus *Bacteroides*

**DOI:** 10.1101/2025.05.07.652691

**Authors:** Archan M. Patel, Howard Ochman

**Affiliations:** Department of Molecular Biosciences, University of Texas at Austin, Austin, Texas 78712, USA

**Author notes:** Correspondence to Howard Ochman.

**Keywords:** gene exchange, speciation, homologous recombination, microbiomes

## Abstract

*Bacteroides* play prominent roles in maintaining gut homeostasis through the fermentation of dietary carbohydrates, the production of metabolites, and the regulation of host immune response. The genus *Bacteroides* comprises several dozen named species, several of which serve as a nexus of gene exchange in gut microbiomes, especially in humans. Because gene exchange will blur the historically defined species boundaries, we assessed the patterns and extent of recombination for nearly 2,000 fully sequenced *Bacteroides* genomes representing 36 named species, and redefined species boundaries based on their capacity to exchange genes. We recognized numerous incongruences between traditional classification schemes, those based on nucleotide sequence identity thresholds, and species defined by gene flow. Almost all named species of sufficient sample size for analysis included genomes that failed to recombine with other genomes that were historically assigned to the same species. Such misclassifications occurred even for cases in which sequence identities surpassed the thresholds regularly used to define species. Conversely, we detected instances in which species previously considered distinct were found to recombine extensively—the most striking of which supported unification of *B. kribbi, B. koreensis*, and *B. ovatus* (which freely recombines with *B. xylanisolvens*). Among the broadly distributed populations of *B. fragilis*, we detected elevated levels of recombination among human-derived strains, although this result is confounded by geographic and sampling biases. Our findings emphasize the utility of a universal species concept based on gene flow, not only for refining bacterial classifications but also for understanding the variation within and across gut microbiomes.

**IMPORTANCE STATEMENT:** *Bacteroides* is a diverse genus comprising some of the most pervasive taxa in the mammalian gut microbiome, with several species playing key roles in the maintenance of gut ecosystem stability. The propensity for *Bacteroides* to exchange genes, both within and among species, has eluded efforts to understand its diversity and to delineate species boundaries, both by traditional classification schemes as well as methods that rely on genomic sequence identity. To circumvent these obstacles and to circumscribe species in a manner that can be uniformly applied across lifeforms, we analyzed all complete genomes assigned to *Bacteroides* and reclassified them based on their capacity for gene flow, uncovering numerous inconsistencies with both recent and historical classifications. Our approach offers a biologically grounded alternative to the use of sequence identity thresholds, offering a better understanding of the diversity and relationships among genomes within the genus *Bacteroides* at large.

## INTRODUCTION

*Bacteroides* are among the most prevalent and abundant bacteria in the human microbiome (1, 2), serving essential functions in degrading polysaccharides (2, 3), modulating immune responses and T-cell development (4, 5), and synthesizing nutrients that sustain the other bacterial species in the gut ecosystem (6). Over 25 species of *Bacteroides* have been isolated from human microbiomes—species that vary greatly in their incidence among individuals and populations, and in their host ranges. *Bacteroides uniformis, B. ovatus*, and *B. xylanisolvens* are the most commonly detected in the human gut (2), along with many other less abundant species that are genetically closely related, though still designated as separate taxonomic groups.

The human microbiome experiences enhanced rates of gene exchange (7, 8), both through the acquisition of new sequences by horizontal gene transfer (HGT) and through the recombinational replacement of homologous sequences (9). And *Bacteroides* species, in particular, have been shown to exhibit elevated levels of both types of gene exchange (10–13). Because homologous recombination targets genes conserved across many strains and species (14), recombining populations can become increasingly genetically similar, thereby complicating the differentiation of species that engage in homologous exchange.

The taxonomy of *Bacteroides* has long been convoluted and subject to frequent revision. Early species were circumscribed based on phenotypic traits [e.g., Gram-negative, non-sporing, rod-shaped, obligately anaerobic (15)], resulting in a genus harboring extreme diversity and spanning a G+C content range of 28–61%, far exceeding the 10% threshold used to distinguish genera at the time (16). To address this heterogeneity, the genus was redefined to include only those strains designated as *Bacteroides fragilis* and its most closely related species (17), whereas the genera *Prevotella* and *Porphyromonas* were established to encompass the excluded species (18, 19). Despite the incorporation of sequencing data, the classification of *Bacteroides* remained imperfect, and even the well-studied bacteria *B. vulgatus* and *B. dorei* were recently committed to a separate genus, *Phocaeicola* (20, 21).

A common approach to bacterial species delimitation involves the application of sequence similarity thresholds, *e.g*., the average nucleotide identity (ANI) of shared genes between strains, to define species membership (22). The Genome Taxonomy Database (GTDB), which endeavors to classify bacteria at all taxonomic levels, uses metrics analogous to ANI to classify genomes based on similarity to clade-specific reference genomes (23). Although these methods are convenient and can be broadly applied, the choice of reference genomes is sometimes arbitrary, and the prescribed ANI thresholds are not based in a biologically guided model of speciation (24, 25).

To assess the extent to which gene exchange has formed or blurred species boundaries in *Bacteroides*, we analyze and classify all available genomes typed to this genus in accord with the Biological Species Concept (26), such that members of a species show evidence of the ability for gene flow, *i.e,* recombination. This feature is regularly used to delineate species of eukaryotes and can be applied across all lifeforms, even viruses (27, 28). We compare this recombination-based classification to the current nomenclature developed for *Bacteroides* as well as to classifications derived from DNA-similarity metrics and assess the degree to which species present in the human microbiome have increased rates of gene exchange.

## MATERIALS AND METHODS

### Analyzed Genomes

All genome sequences assigned to the genus *Bacteroides*, available as of April 2024, were downloaded from the NCBI RefSeq database (www.ncbi.nlm.nih.gov/refseq), and are available via the accession numbers listed in Table S1. Genomes were selected to ensure that analyses excluded redundant, incomplete, or metagenomically derived genomes. Genome assemblies were evaluated with CheckM (29), and those that were <99% complete or that contained >1% contamination were removed. This filtering process flagged 1,271 genomes for removal and yielded a final dataset of 1,926 genomes representing 36 nominal species of *Bacteroides*, as follows: *B. bouchesdurhonensis* (*n* = 3), *B. caccae* (*n* = 70), *B. cellulosilyticus* (*n* = 45), *B. clarus* (*n* = 9), *B. congonensis* (*n* = 1), *B. cutis* (*n* = 2), *B. eggerthii* (*n* = 46), *B. faecichinchillae* (*n* = 1), *B. faecis* (*n* = 25), *B. faecium* (*n* = 1), B*. fluxus* (*n* = 3), *B. fragilis* (*n* = 354), *B. gallinaceum* (*n* = 9), *B. graminisolvens* (*n* = 2), *B. helcogenes* (*n* = 1), *B. hominis* (*n* = 11), *B. humanifaecis* (*n* = 2), *B. intestinalis* (*n* = 68), *B. koreensis* (*n* = 2), *B. kribbi* (*n* = 1), *B. luhongzhouii* (*n* = 1), *B. mediterraneensis* (*n* = 3), *B. nordii* (*n* = 14), *B. oleiciplenus* (*n* = 3), *B. ovatus* (*n* = 252), *B. parvus* (*n* = 1), *B. pyogenes* (*n* = 21), *B. reticulotermitis* (*n* = 1), *B. rhinocerotis* (*n* = 1), *B. salyersiae* (*n* = 50), *B. stercorirosoris* (*n* = 1), *B. stercoris* (*n* = 61), *B. thetaiotaomicron* (*n* = 288), *B. uniformis* (*n* = 297), *B. xylanisolvens* (*n* = 161), *Candidatus B. intestinigallinarum* (*n* = 3), and 112 genomes classified as *Bacteroides* but not assigned to any species (Table S1).

### Taxonomic assignment of genomes

To test the veracity of the NCBI species assignments, we subjected the *Bacteroides* dataset to three sequence-based methods that have been employed to assign genomes to species:

1. <u>Average Nucleotide Identity</u> (ANI) quantifies the genetic similarity between two genomes by calculating the mean nucleotide sequence identity across all orthologous genes. We used the “many-to-many” option in *FastANI* v1.33 (30) to compute the ANI between every pair of genomes in our dataset. ANI values were evaluated in relation to the nominal species designations in the NCBI and the classification scheme offered by the GTDB (23), which is itself based on a normalized ANI metric.

2. *<u>ConSpeciFix</u>* is a computational pipeline that detects recombinant alleles in alignments of the orthologous genes shared by a set of genomes (*i.e*., the core genome). We define orthologous genes as those exhibiting >70% identity in pairwise USEARCH comparisons and differing by no more than 20% in length. To build core genome alignments, we required that single-copy orthologs be present in at least 85% of the strains considered. In *ConSpeciFix*, species boundaries are delineated based on the capacity for gene exchange, such that lineages that recombine are considered members of the same biological species. We utilized *ConSpeciFix* v1.3.0 (31) to estimate the ratio of homoplastic alleles (*h;* those not inherited vertically) to non-homoplastic (*m;* those inherited vertically or due to new mutations) alleles from randomly sampled subsets of *Bacteroides* genomes. The inclusion of a non-recombining genome in the analyzed subset of strains results in a reduction in *h/m* ratios, indicating its status as a separate biological species. The *ConSpeciFix* output graphs were visually inspected to confirm the recombination status of each group.

To assess which genomes could be consolidated into a single biological species, we first grouped and analyzed the genomes from each of the 11 NCBI-designated species that contained ≥30 genomes (termed “focal” species) to determine which, if any, strains were not recombining with the others classified as the same species. To detect non-recombining strains, we extracted random groups of 30 genomes from the total number available for each of the 11 focal species and ran *ConSpeciFix* on each group. We elected to use groups of 30 strains instead of analyzing all genomes of a focal species simultaneously because smaller sample sizes both provide better granularity for detecting the influence of individual strains on *h/m* values and make analyses more computationally manageable. We determined the number of 30-strain groups necessary to fully envelop each focal species by performing 10,000 simulations: for each simulation, we randomly sampled groups of 30 strains and noted how many groups were needed until every strain was represented at least once. The average number of groups needed across all simulations was used as the final number of 30-strain groups for each species. Non-recombining genomes were identified and removed from each group, leaving only those considered to represent a single biological species.

Next, we determined species boundaries within *Bacteroides* by examining changes in the *h/m* ratios after integrating genomes from another NCBI species (“test lineages”) with each focal lineage. The test lineages included in these analyses represented those genomes classified to a distinct NCBI species and having the highest and lowest ANI in relation to a selected focal species. For cases in which test lineages were also members of one of the focal species, we selected the test lineages from those determined to comprise the biological species of its assigned focal species (*i.e.*, only genomes that were confirmed to recombine); in all other cases, we selected test lineages from among all genomes assigned by the NCBI to a particular species. For NCBI species with more than 30 non-redundant genomes, we restricted our analysis to 30 randomly selected genomes to enhance computational efficiency.

3. *<u>PopCOGenT</u>* is an alternate approach to species delineation that is based on the premise that regions introduced by homologous exchange and/or HGT will display fewer substitutions than those inherited vertically. *PopCOGenT* flags pairs of genomes as recombinant—and thus belonging to the same species—if they have longer and more prevalent identical regions than would be expected by mutations alone (32). We applied *PopCOGenT* to the group of 51 *Bacteroides* genomes, manually curated to include only non-redundant genomes that were labeled differently across the various classification schemes.

### Phylogenetic Analysis

A maximum-likelihood phylogeny based on the 51 *Bacteroides* genomes was generated in the UCBG2 pipeline (33), using a set of 81 universally conserved, single-copy genes, as identified with Prodigal (34) and HMMER (35). Extracted gene sequences were aligned with MAFFT (36), and the phylogenetic tree was generated with FastTree (37) using the generalized time-reversible model of nucleotide evolution.

## RESULTS

Below, we evaluate the species status of the 11 nominal species of *Bacteroides* for which at least 30 full genome sequences were available in the NCBI RefSeq database (www.ncbi.nlm.nih.gov/refseq/). We compared and classified the genomes of these species relative to one another and to 25 more sparsely sampled *Bacteroides* species for which one or more full genome sequences have been resolved.

### Bacteroides caccae

Based on *ConSpeciFix*, all genomes designated as *B. caccae* in the NCBI RefSeq database are members of a single recombining species, with an *h/m* value—the ratio of homoplastic to ancestrally-derived polymorphisms—that plateaus at 1.7 (Figure 1A). Strains classified as *B. caccae* are very similar at the DNA level, with an average nucleotide identity (ANI) of 98.3% between the most divergent conspecifics and an average pairwise ANI of 99.1% among all strains.

**Figure 1.**
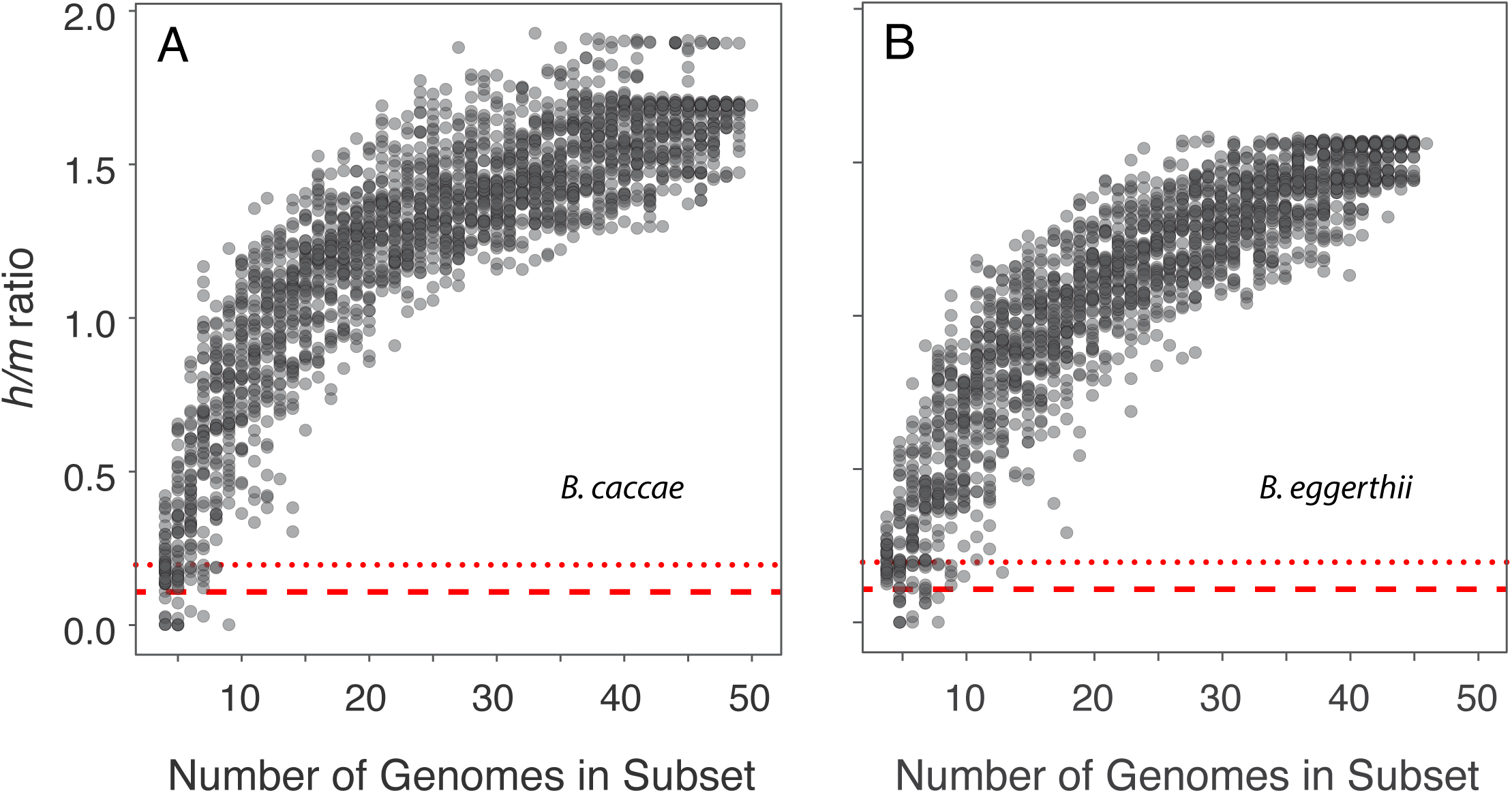
Recombinational analysis of (A) *Bacteroides caccae* and (B) *Bacteroides eggerthii.* Ratio of homoplastic (*h*) to non-homoplastic (*m*) polymorphisms for randomly sampled subsets of the NCBI-designated genomes assigned to the designated species. The red dashed line indicates the average *h/m* ratio expected if homoplasies arise exclusively from convergent mutations, and the red dotted line indicates the maximum *h/m* ratio expected if homoplasies arise exclusively from convergent mutations. The *h/m* values specified by the two red lines were estimated by simulating sequence evolution without recombination across 97 species (Bobay and Ochman, 2017).

### Bacteroides cellulosilyticus

Unlike *B. caccae*, certain strains designated as *B. cellulosilyticus* do not participate in gene exchange with the others and constitute a distinct species. Analyses of randomly sampled subsets of *B. cellulosilyticus* genomes revealed that nine strains, when included in the datasets, consistently lowered *h/m* values (Table S2, Figure 2A). When these nine strains are excluded from the analyses, *h/m* values increased from 0.6–0.8 (depending on which of the nine strains were included) to 1.1, and *ConSpeciFix* regards all remaining strains as representing a single recombining species (Figure 2B).

**Figure 2.**
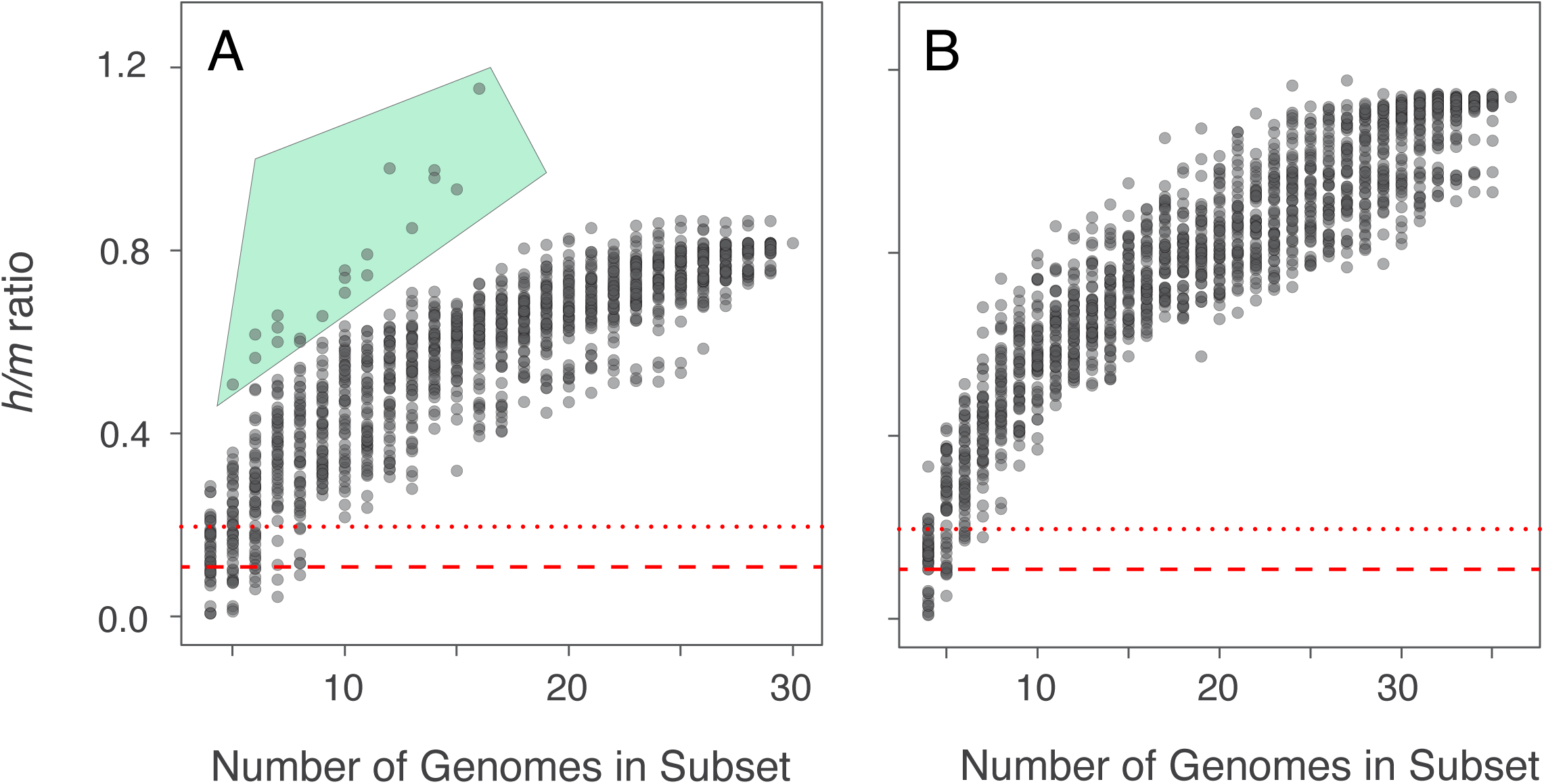
Recombinational analysis of *Bacteroides cellulosilyticus*. **(A)** Ratio of homoplastic (*h*) to non-homoplastic (*m*) polymorphisms for randomly sampled subsets of NCBI-designated *B. cellulosilyticus* genomes. Subsampled sets of genomes with the highest *h/m* values (those enclosed within the green polygon) all lack one or more representatives of a specific set of nine genomes. **(B)** Analysis of subsets of *B. cellulosilyticus* genomes after excluding the nine aforementioned genomes. The two red lines indicate *h/m* values expected when homoplasies arise exclusively from convergent mutations as described in the legend to Figure 1.

The Genome Taxonomy Database (GTDB), using a scaled ANI-based method, also regards these nine strains as a separate species, grouping them under the appellation *Bacteroides sp900552405*. After excluding these strains, the average pairwise ANI among *B. cellulosilyticus* genomes is 98.1%, with an ANI of 97.1% between the most divergent pair. The nine strains designated as *Bacteroides sp900552405* are themselves fairly closely related, with a mean ANI of 99.6%, and as a group, they average 93.9% ANI to the *B. cellulosilyticus* genomes that constitute a recombining species.

### Bacteroides eggerthii

All of the 46 genomes designated as *B. eggerthii* in the RefSeq database belong to a single recombining species, with an *h/m* ratio plateauing at 1.6 (Figure 1B). Strains of *B. eggerthii* have an average pairwise ANI of 99.0% and an ANI of 98.2% between the most divergent conspecifics.

### Bacteroides fragilis

*ConSpeciFix* analyses indicate that the genomes designated as *B. fragilis* in the RefSeq database constitute multiple species. Inclusion of 98 strains (Table S3) consistently decreased *h/m* values, and their removal from analyses yielded a single recombining species (Figure 4A, B). The GTDB classifies 97 of these genomes as *B. hominis* and one as *B. fragilis_*B. The average pairwise ANI among the 256 recombining strains of *B. fragilis* genomes is 98.9%, with an ANI of 98.2% between the most divergent conspecifics. The excluded strains average an ANI of 98.2% with one another and are highly divergent from *B. fragilis,* with an average ANI of 87.4% from the genomes that constitute a recombining species.

### Bacteroides intestinalis

Six strains designated as *B. intestinalis* consistently lowered *h/m* values when included in the analyzed subsets of genomes (Table S4). In analyses that tested each of these strains with those of other species of *Bacteroides*, one of these strains (GCF_001578635.1) was found to recombine with *B. cellulosilyticus*, and the GTDB also reclassifies it as such. The five other non-recombining strains have also been reclassified by the GTDB—four as *B. intestinalis_A* and one as *B. sp900556215*.

After excluding these six strains from analyses, all remaining strains (*n* = 62) constitute a single recombining species (Figure 3A). In contrast, the GTDB reclassifies 27 of the 62 strains that *ConSpeciFix* established as a single species as *Phocaeicola sp948566455*. However, removing those 27 strains from analyses did not appreciably alter *h/m* values, either overall or at the corresponding subsample sizes (Figure 3B), indicating that they do not form a separate species and should all be considered *Bacteroides intestinalis.* These 62 strains (including those regarded as *Phocaeicola sp948566455*) have an average pairwise ANI of 98.3%, with the most divergent pair displaying 97.1% identity. Notably, the average pairwise ANI does not change when the 27 strains classified as *Phocaeicola sp948566455* by the GTDB were excluded, establishing that the GTDB-reclassified strains are subsumed within the variation of *B. intestinalis* and should continue being considered members of that species. The six non-recombining strains have an average ANI of 94.7% to one another and an average ANI of 93.3% from the genomes that constitute a recombining species.

**Figure 3.**
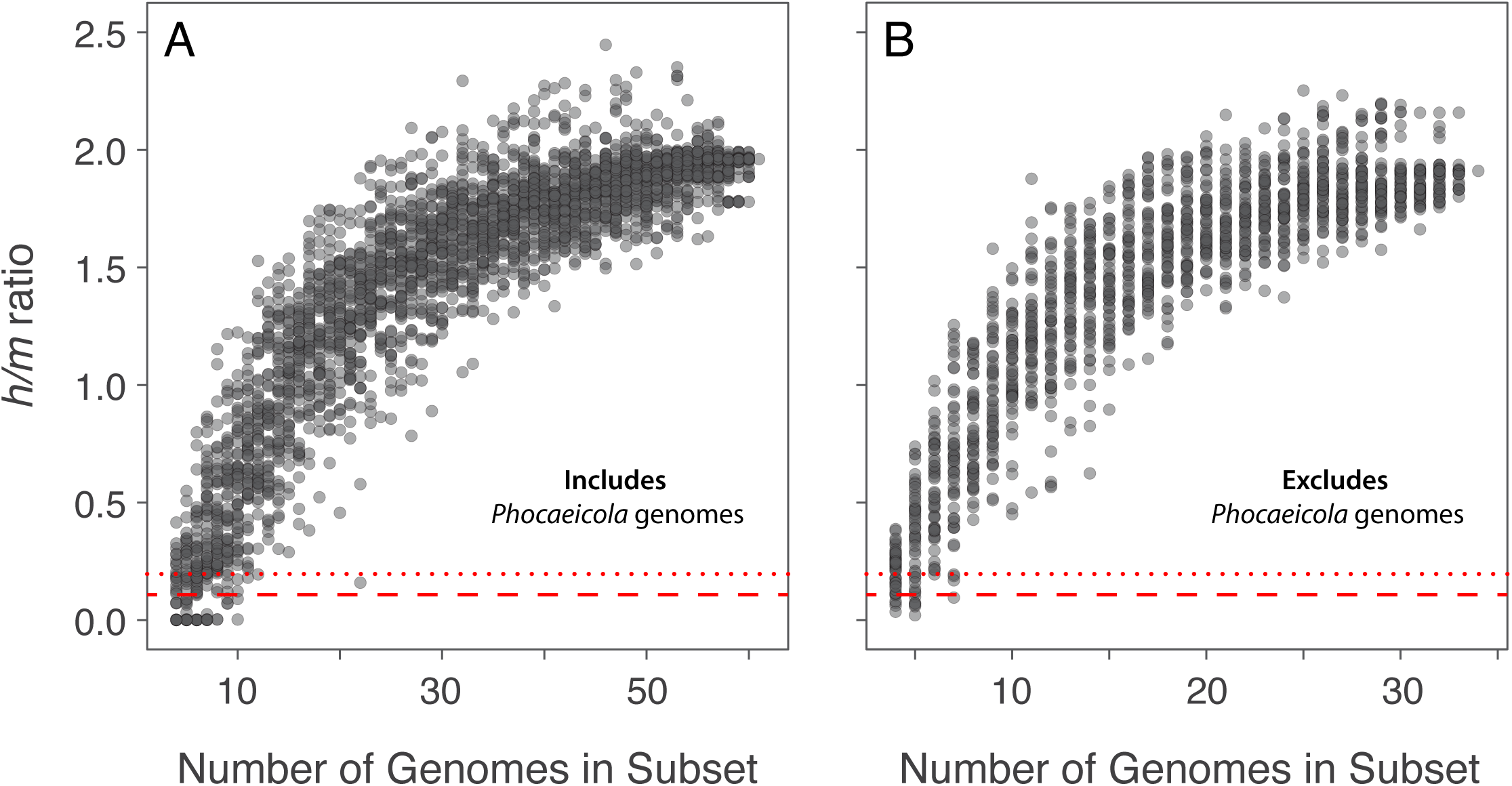
Recombinational analysis of *Bacteroides intestinalis*. **(A)** Ratio of homoplastic (*h*) to non-homoplastic (*m*) polymorphisms for randomly sampled subsets of 61 NCBI-designated *B. intestinalis* genomes, which includes genomes later assigned to *Phocaeicola sp948566455* by the GTDB. **(B)** *h/m* ratios after excluding strains regarded as *Phocaeicola sp948566455* from analyses of *B. intestinalis* genomes. That panels **A** and **B** are virtually identical, regardless of whether the GTDB-classified strains of *Phocaeicola sp948566455* are included in analyses, indicates that all strains are members of a single recombining species. The two red lines indicate *h/m* values expected when homoplasies arise exclusively from convergent mutations.

### Bacteroides ovatus

Of the 252 *B. ovatus* strains available in the RefSeq database, *ConSpeciFix* regards all but one as members of a single recombining species (Figure 4C, D). The single strain (GCF_021410395.1) does not engage in gene exchange with the others and has been reclassified by the GTDB as *B. sp902362375* (Table S5). The recombining strains of *B. ovatus* display an average pairwise ANI of 97.4% and an ANI of 95.8% between the most divergent pair. The non-recombining strain averages an ANI of 92.8% to the *B. ovatus* genomes that constitute a single species.

### Bacteroides salyersiae

Analyses of the 50 strains designated as *B. salyersiae* reveal that three did not recombine with the others (Table S6, Figure 4E). Upon exclusion of these three non-recombining strains, *h/m* ratios, when viewed across subsample sizes, do not produce the smooth plateau expected for a single recombining population. Instead, *h/m* values are diffuse across the range of subsample sizes, although no specific strains consistently appear in subsets with higher or lower *h/m* values. On account of this inability to detect strains that consistently modify *h/m* ratios, we extracted the core set of genes from each genome using *CoreCruncher,* which can sometimes improve the alignments built from large sets of bacterial genomes (Harris et al. 2020). Analysis of the refined core gene set generated an *h/m* plot that mirrors what is typically observed for a single recombining species (Figure 4F).

After excluding the non-recombining strains, the remaining *B. salyersiae* genomes show high sequence similarity, displaying an average pairwise ANI of 99.4%, with the most divergent pair having 98.8% identity. The three non-recombining strains, which in this instance, the GTDB does not view as a separate species, have an average pairwise ANI of 97.3% with one another and an average ANI of 97.9% to the recombining strains (Table S6).

### Bacteroides stercoris

Two of the 61 strains designated as *B. stercoris* do not recombine with the others (Figure 4G), and the remaining 59 strains constitute a single recombining species, with an *h/m* value plateauing at 1.5 (Figure 4H). The two non-recombining strains— GCF_900624775.1 and GCF_900624875.1—have been reclassified by the GTDB as *B. eggerthii* and *B. clarus,* respectively (Table S7). *ConSpeciFix* supports the reclassification of GCF_900624775.1 as *B. eggerthii* in that it recombines with other *B. eggerthii* strains, but the veracity of the second reclassification cannot be assessed because too few additional *B. clarus* genomes are available for comparison.

Upon exclusion of the two non-recombining strains, the average pairwise ANI among *B. stercoris* genomes is 98.2%, with the most divergent pair of strains having an ANI of 97.0%. The two non-recombining strains have a pairwise ANI of 86.8% and are highly diverged from *B. stercoris,* with an average pairwise ANI of 86.5%.

### Bacteroides thetaiotaomicron

Only seven of the 288 NCBI-designated *B. thetaiotaomicron* genomes do not engage in gene exchange with the others (Table S8, Figure 4I), and after excluding these seven genomes from the analysis, all remaining strains of *B. thetaiotaomicron* constitute a single recombining species, with an *h/m* value plateauing at 2.35 (Figure 4J). The 281 *B. thetaiotaomicron* strains forming a single recombining species are closely related at the nucleotide level, with an average pairwise ANI of 98.1% among strains, and an ANI of 96.3% between the most divergent conspecifics. The seven non-recombining strains, which are not classified as distinct species by the GTDB, have an average ANI of 98.3% with one another and are only slightly more distantly related (ANI = 97.9%) to the *B. thetaiotaomicron* strains that constitute a single recombining species.

### Bacteroides uniformis

Of the 297 strains designated as *B. uniformis* in the RefSeq database, *ConSpeciFix* identified 24 that consistently decreased *h/m* ratios when included in the analyzed subsets (Table S9, Figure 4K). Upon the removal of these genomes from the analysis, the remaining 273 strains form a single recombining species, with an *h/m* ratio that plateaus at 1.8 (Figure 4L).

The 273 strains of *B. uniformis* that constitute a recombining species exhibit an average pairwise ANI of 98.2%, with an ANI of 96.7% between the most distant pair. The 24 non-recombining strains have an average pairwise ANI of 98.7% with one another and an average pairwise ANI of 96.1% with the recombining *B. uniformis* strains. Similar to the situation in *B. thetaiotaomicron*, the non-recombining strains of *B. uniformis* are not viewed as distinct species in the GTDB on account of their high ANI to the cohort of recombining strains.

### Bacteroides xylanisolvens

Ten of the 161 genomes designated as *B. xylanisolvens* do not engage in gene exchange with the others (Figure 4M). Excluding these ten strains from the analysis leaves a single recombining species with an *h/m* ratio plateauing at 2.2 (Figure 4N). The GTDB considers nine of the excluded strains, along with a strain previously designated as *B. ovatus* (see above), as *Bacteroides sp902362375*, and the remaining strain is classified as *B. luhongzhouii* (Table S10).

The 151 genomes of *B. xylanisolvens* that form a single recombining species display an average pairwise ANI of 97.5%, with an ANI of 96.1% between the most divergent pair. The non-recombining strains have an average pairwise ANI of 90.2% to the strains that constitute *B. xylanisolvens* but are themselves closely related, with a mean pairwise ANI of 97.9%.

### Recombination barriers between *Bacteroides* lineages

To further delineate species boundaries within *Bacteroides*, we assessed whether any of the 11 species considered above (and now defined based on recombination among members) engage in gene exchange with one another or with lineages from any of 25 other NCBI-designated *Bacteroides* species. We repeated our analyses on each of these 11 focal species after including one or more genomes from another species and evaluated the resulting *h/m* values for shifts indicative of the ability, or the lack of ability, of the newly included genome(s) to recombine with the focal species. Whereas the majority of strains typed to different named species show no evidence of gene exchange with one another, there were obvious cases (discussed below) in which members of an alternate species recombined with the tested focal species, necessitating merging and re-classification.

### Bacteroides ovatus subsumes Bacteroides koreensis

*ConSpeciFix* analyses display no reduction in *h/m* ratios when the two available *B. koreensis* genomes were incorporated into subsets of *B. ovatus* genomes that were previously determined to comprise a single biological species. The ANI between *B. koreensis* and *B. ovatus* genomes (96.3% maximal divergence) fall within the range observed for *B. ovatus* genomes. 95.8% maximal divergence). Only one of the two *B. koreensis* genomes is listed in the GTDB database, where it is reclassified as *B. ovatus* (Table S11).

### Bacteroides ovatus subsumes Bacteroides kribbi

*B. kribbi* and *B. ovatus* constitute a single recombining species, as evidenced by the persistent smooth curve of *h/m* ratios when the single genome assigned to *B. kribbi* was included with *B. ovatus* subsets. In fact, the results are unchanged when *B. ovatus, B. koreensis* and *B. kribbi* are all included in the same analysis, indicating that these three nominal species should be deemed the same biological species (Figure 5A). *B. ovatus* and *B. kribbi* have a high average pairwise ANI of 97.6% and an ANI of 96.6% between the most divergent pair. This high degree of sequence similarity prompted the GTDB to reclassify the strain designated as *B. kribbi* to *B. ovatus* (Table S12).

**Figure 4.**
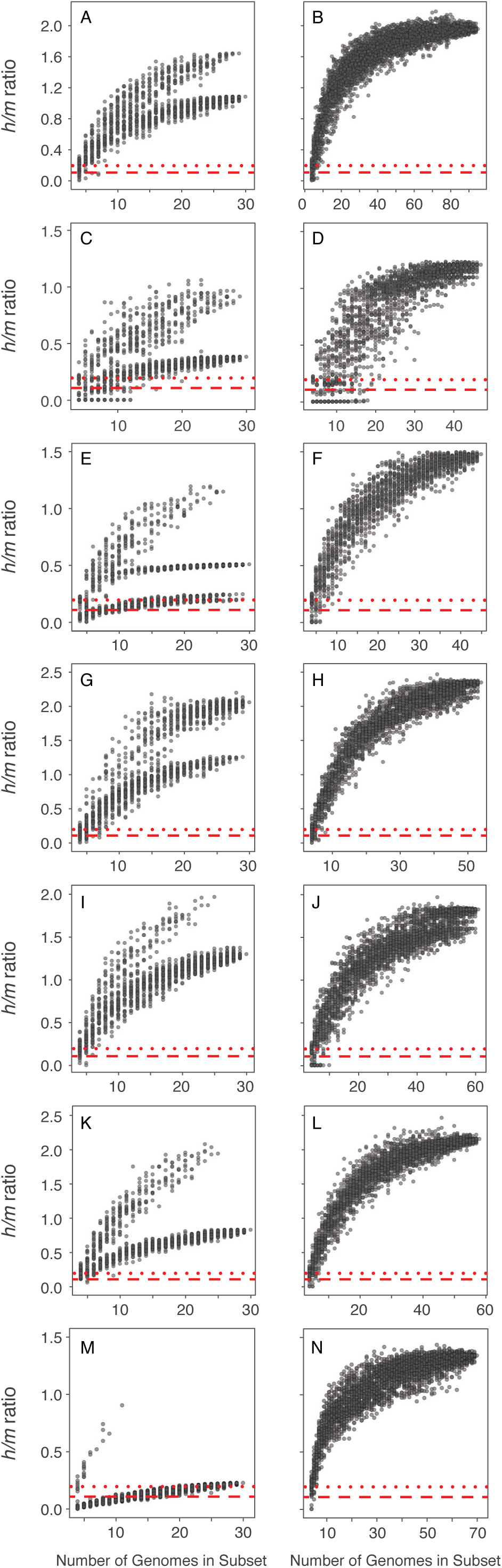
Recombinational analysis of NCBI-designated species of *Bacteroides.* Ratio of homoplastic (*h*) to non-homoplastic (*m*) polymorphisms for randomly sampled subsets of genomes from NCBI-designated species of *Bacteroides*. For each species shown, one or more genomes assigned to the particular NCBI-designated species did not recombine with other genomes assigned to that species resulting in the formation of multiple plateauing curves when analyzed with *ConSpeciFix* (left-hand panels). In each case, the subsequent identification and exclusion of the non-recombining genomes from analyses yielded a set of genomes comprising a single recombining species with *h/m* ratios that plateau at a higher value (right-hand panels). **(A)** *B. fragilis*, **(B)** *B. fragilis* after exclusion of 107 non-recombining genomes; **(C)** *B. ovatus,* **(D)** *B. ovatus* after exclusion of the single non-recombining genome; **(E)** *B. salyersiae,* **(F)** *B. salyersiae* after exclusion of 3 non-recombining genomes; **(G)** *B. stercoris,* **(H)** *B. stercoris* after exclusion of two non-recombining genomes; **(I)** *B. thetaiotaomicron,* **(J)** *B. thetaiotaomicron* after exclusion of non-recombining genomes; **(K)** *B. uniformis,* **(L)** *B. uniformis* after exclusion of non-recombining genomes; **(M)** *B. xylanisolvens,* **(N)** *B. xylanisolvens* after exclusion of non-recombining genomes. The two red lines indicate *h/m* values expected when homoplasies arise exclusively from convergent mutations.

**Figure 5.**
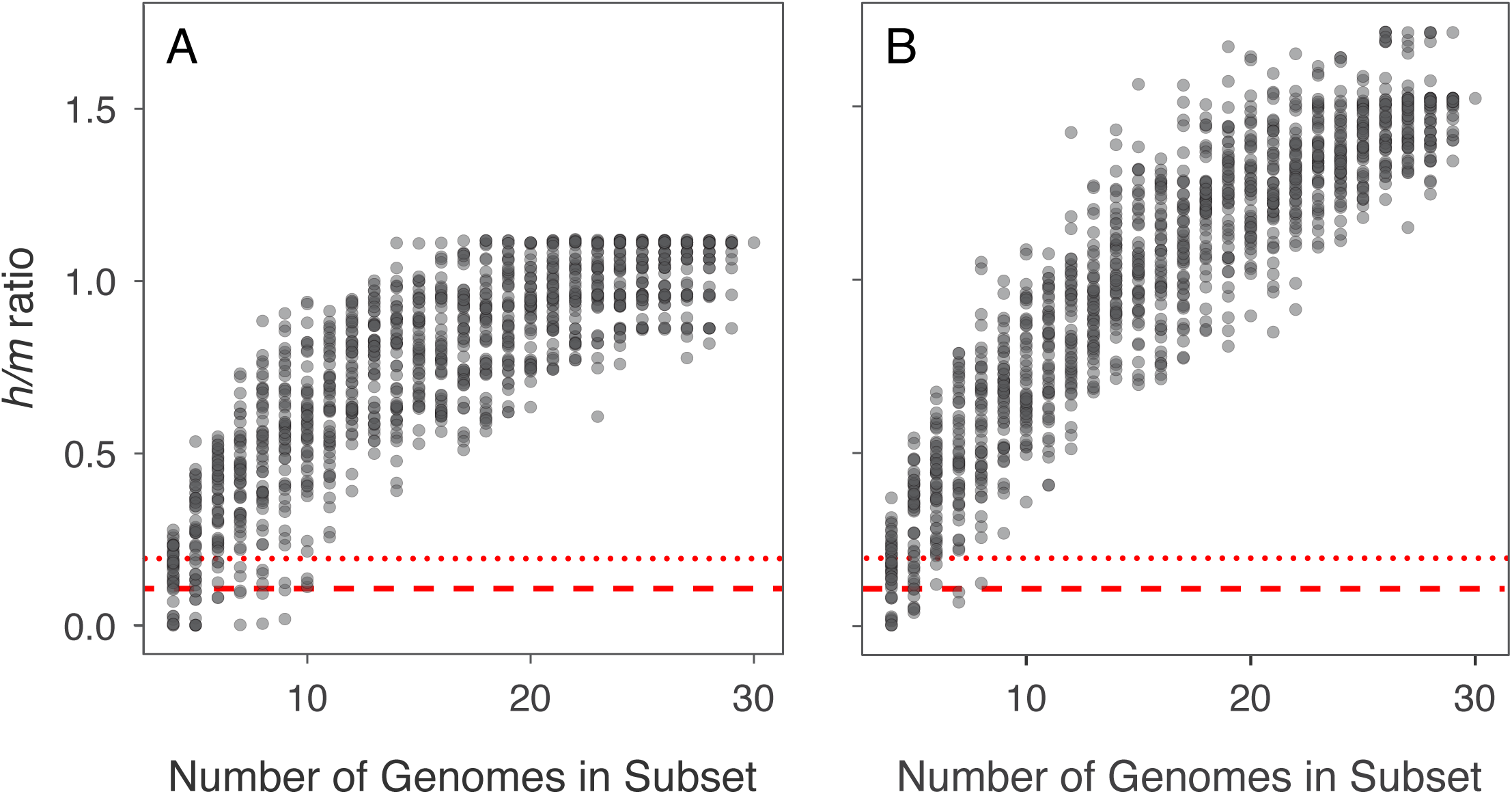
Recombination between *Bacteroides ovatus* and other nominal species of *Bacteroides.* **(A)** Ratio of homoplastic (*h*) to non-homoplastic (*m*) polymorphisms for randomly sampled subsets of *Bacteroides ovatus* and all available *B. koreensis* (*n* = 2) and *B. kribbi* genomes (*n* = 1). The single plateauing curve indicates that genomes assigned to *B. ovatus*, *B. koreensis,* and *B. kribbi* constitute a single recombining species. **(B)** *h/m* values for randomly sampled subsets of *B. ovatus* genomes when including the most closely related and most distantly related *B. xylanisolvens* genomes. Again, the single plateauing curve indicates that genomes assigned to *B. ovatus* and *B. xylanisolvens* represent a single recombining species. The two red lines indicate *h/m* values expected when homoplasies arise exclusively from convergent mutations.

### Bacteroides ovatus and Bacteroides xylanisolvens

Based on *ConSpeciFix*, strains belonging to *B. ovatus* and *B. xylanisolvens* constitute a single biological species, despite their low nucleotide similarity: average pairwise ANI between *B. ovatus* and *B. xylanisolvens* genomes is 92.9%, and only 91.6% among the most divergent pair. Plots of *h/m* ratios maintain strong signals of gene flow when either the most closely related or the most divergent strains of *B. xylanisolvens* relative to *B. ovatus* are included (Figure 5B) and also when equal numbers *B. ovatus* and *B. xylanisolvens* strains are randomly sampled. In contrast to our conclusion that *B. ovatus* and *B. xylanisolvens* constitute a single species, the GTDB views them uniformly as representing two distinct species.

Although *B. ovatus*, *B. kribbi,* and *B. koreensis* recombine with one another (Figure 5A), as do *B. ovatus* and *B. xylanisolvens* (Figure 5B), there is no signal of recombination between *B. xylanisolvens* and either *B. koreensis* or *B. kribbi.* That *B. xylanisolvens* recombines with *B. ovatus* but not with *B. kribbi* and *B. koreensis* is surprising given that it has approximately the same level of nucleotide similarity with all three species.

### Bacteroides uniformis subsumes Bacteroides humanifaecis

Both of the NCBI-designated *B. humanifaecis* strains available for analysis engage in gene flow with *B. uniformis* (Figure S1). The average pairwise ANI among *B. uniformis* and *B. humanifaecis* strains is 96.0%, and consistent with the results from *ConSpeciFix,* the GTDB reclassifies both of these *B. humanifaecis* strains as *B. uniformis* (Table S13).

## DISCUSSION

*Bacteroides* are common constituents of mammalian gut microbiomes (38), and by applying a recombination-based approach, we established the species boundaries and delineations of all available genome sequences assigned to this genus. Although many current *Bacteroides* species classifications were upheld, numerous genomes and species assignments defy previous nomenclature and/or classification based on DNA similarity indices.

Of the 11 well-sampled nominal species that we tested (i.e., our “focal” species), only one (*B. caccae*) aligned perfectly with current designations, such that all genomes classified to this species in the RefSeq database are members of a single biological species, and no genomes classified as a different species need to be incorporated into this species. Many of the other nominal species incurred minor changes: for example, all of the 46 genomes classified as *B. eggerthii* form a single species; however, one strain of *B. stercoris* needs to be reclassified as a member of *B. eggerthii*. Additionally, there are several cases in which one or a few genomes were incorrectly assigned to a particular species. Only 1 of the 252 *B. ovatus* genomes, 2 of the 61 *B. stercoris* genomes, and 3 of the 50 *B. salyersiae* genomes were misclassified.

Among the 11 focal species, *B. fragilis* contains the largest proportion of genomes that required reclassification due to the absence of gene flow with the other strains assigned to that species. Historically, *B. fragilis* has been separated into two groups, Division I and Division II, based on the presence of the antimicrobial resistance gene *cepA* (39, 40). Other genetic-based methods have consistently distinguished the two groups (39, 41–43), prompting proposals to recognize Division II as a distinct species (44). Various species-delineation approaches classify *B. fragilis* Division II strains with *B. hominis*—a species formally proposed in 2021 (45)—however, the reclassification has not been adopted universally (46). Our analyses fully support their reassignment: almost 30% of the more than 300 genomes originally classified as *B. fragilis* exchange genes with one another and with strains classified as *B. hominis* but show no evidence of exchange with the recombining *B. fragilis* population.

Several other species refinements based on the interruption of gene flow are congruent with the ANI-threshold-based classifications imposed by the GTDB. For example, our analyses detected non-recombining strains within NCBI-designated *B. cellulosilyticus, B. ovatus*, *B. stercoris* and *B. xylanisolvens*, and in each of these cases, the GTDB also advised reassignment of the same strains to alternate species. That the GTDB and *ConSpeciFix* earmark the same strains is an indication that distinct biological species often abide the 95% ANI threshold and that 5% sequence divergence can be sufficient to disrupt homologous exchange. In addition to the separation of genomes based both on recombination and the application of a 95% ANI threshold, these same criteria recommended consolidation of separate nominal species (*e.g*., *B. ovatus, B. koreensis* and *B. kribbi*) into a single biological species.

However, strict application of a 95% ANI-threshold for species delineation is undefendable: in many cases, we detected no signal of gene exchange for certain strains despite their having >95% ANI with the recombining members of the species (e.g., *B. parvus* and *B. rhinocerotis* with *B. uniformis*). Conversely, we also identified genomes sharing <95% ANI (and designated as different species by the NCBI and the GTDB) that recombine with one another. For example, *B. xylanisolvens* and *B. ovatus* have an average pairwise ANI of 92.9% but offer clear evidence of recombination and should be considered a single species. Recombination efficiency drops as strains share diminishing nucleotide identity, but the specific threshold at which recombination breaks down varies among bacterial species (27, 47–49), thereby confounding the universal application of any single ANI threshold to delineate bacterial species.

Comparing the species classification offered by the GTDB with those based on gene flow not only highlights the inconsistencies arising from application of ANI thresholds but also reveals instances in which species recognition has relied on historical convention and/or methodological constraints as opposed to biological criteria. For example, the GTDB regards a set of 27 of the 61 *B. intestinalis* strains as a separate species (dubbed *Phocaeicola sp948566455*); however, these strains recombine to the same degree with one another and with genomes classified as *Bacteroides intestinalis*. Curiously, the type strains for *Phocaeicola sp.* and *B. intestinalis* in the GTDB database have >95% ANI despite being classified as separate species. This unwarranted differentiation of species reflects vagaries of the GTDB protocol, which elects to retain distinct classifications for species representatives that share 95–97% ANI and merges them only if they share >97% ANI (50). Because the *Phocaeicola sp.* type strain was designated as such in the literature, the GTDB’s ANI-based approach enforces this distinct classification.

*PopCOGenT* offers an alternate recombination-based approach for delimiting species boundaries by grouping genomes that share longer and more frequent stretches of identical regions than expected solely from the accumulation of mutations (32). Both *ConSpeciFix* and *PopCOGenT* adhere to the precepts of the Biological Species Concept, such that genomes connected through gene exchange are considered members of the same species; however, the methods differ in criteria for identifying recombination and, very often, species boundaries. *ConSpeciFix* assesses recombination in the set of genes shared by all genomes, whereas *PopCOGenT* searches for similarities across the entire genome, including those accessory regions present only in a subset of strains. By including sporadically distributed regions, *PopCOGenT* can group distant genomes that engaged in recent lateral gene transfer as well as those that participate in homologous exchange and will consequently expand species boundaries. This feature is evident in the *Bacteroides* species designations resolved by the GTDB, *ConSpeciFix,* and *PopCOGenT* (Figure 6): *PopCOGenT*, in contrast to the other classification schemes, considers vastly divergent strains (*i.e*., those bearing the same *PopCOGenT* number) to comprise a single biological species.

**Figure 6.**
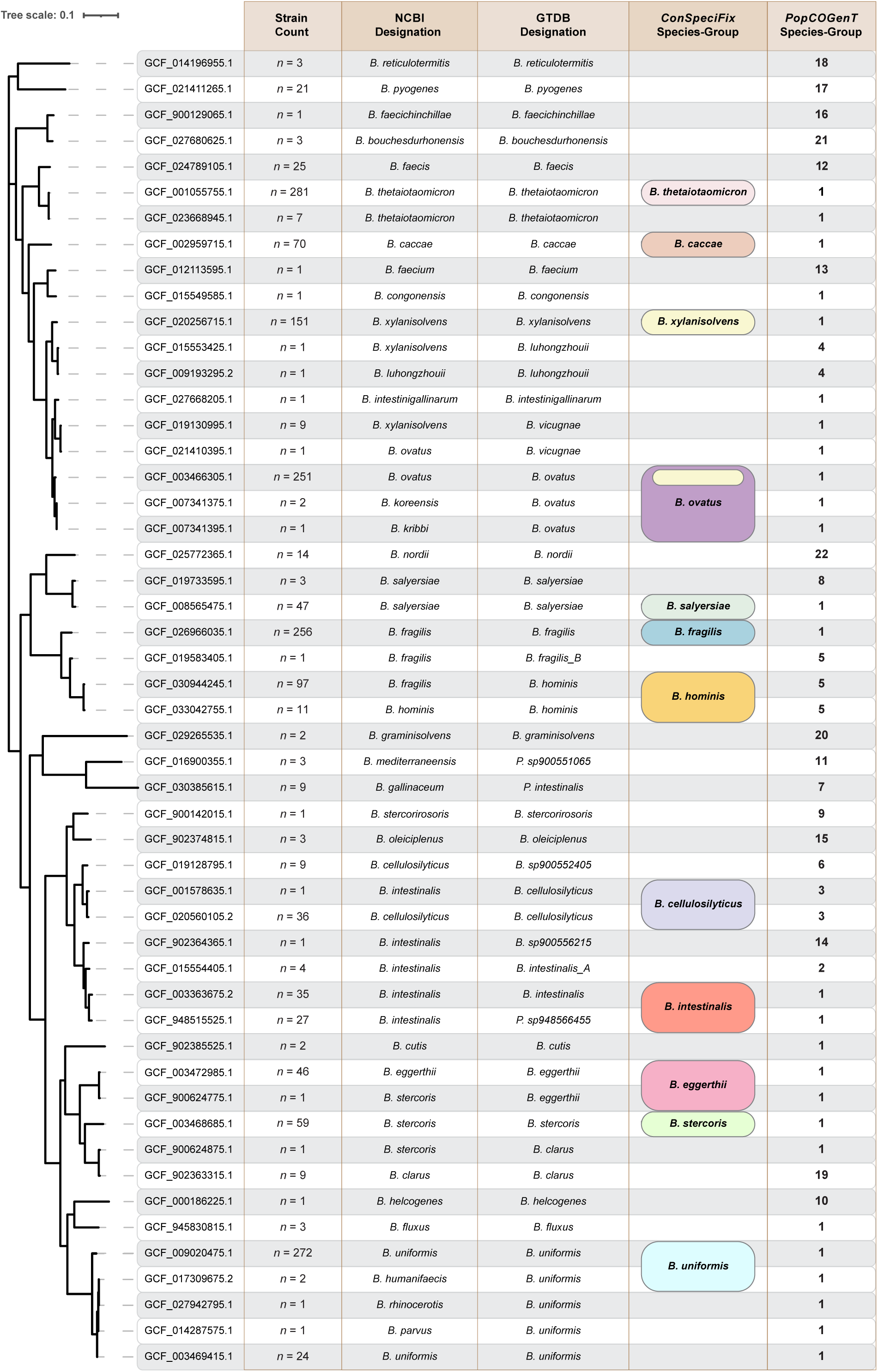
Species assignment in the genus *Bacteroides*. Maximum-likelihood phylogeny and species-level classifications of representative *Bacteroides* genomes based on different classification schemes. Genomes were selected both to depict the diversity within the genus and to highlight discrepancies between the classification systems. Strain Count column includes the number of genomes that exhibit the identical nomenclatures and species-group assignments across classification systems, with the accessions, sampled randomly from each group, used to facilitate *PopCOGenT* analysis. In the *ConSpeciFix* column, the lack of a latin binome indicates that the corresponding genome(s) do not recombine with members of the 11 focal species; in the *PopCOGenT* column, all genomes with the same number are viewed as members of the same species-group.

Several groups have reported enhanced rates of gene exchange among strains that inhabit the human microbiome (7, 8, 51). Many of these studies infer gene transfer events based on homology (7, 8)—an approach that works well when genes from distantly related sources are introduced into a genome but often fails to detect recombinational exchange among genes shared by members of the same species (14). Moreover, the detected association between exchange rates and incidence in the human microbiome may possibly result from sampling biases, since more human-associated species have been sequenced (52).

A recent study that focused specifically on the levels of homologous exchange occurring in the human microbiome reported high levels of recombination among human-associated *Bacteroides* (12). Given the broad assemblage of species considered in that study, we can directly compare one of their metrics of recombination (*T_mrca_/T_mosaic_*) to the analogous *h/m* values calculated in our analyses for seven of our focal species. We detect no significant correlation between these two measures of recombination (Kendall’s Tau = 0.048; *p* > 0.9; *n* = 7) (Table S14), an incongruence possibly due to the inclusion of non-human-derived strains in some of the species we evaluated.

We tested this issue in two ways: first, we eliminated all strains isolated from non-human hosts prior to performing our analysis, but, again, there was no association between the levels of recombination detected in the two studies (Kendall’s Tau = 0.1; *p* = 0.75; *n* = 7). Next, we compared recombination rates in the human-derived *B. fragilis* to a group of *B. fragilis* strains derived exclusively from non-human hosts, which included isolates from pigs, chickens, and orangutans. The human-derived *B. fragilis* strains exhibit elevated rates of recombination compared to the non-human-derived strains (Figure S2), a consequence of either an actual increase in recombination rates or that human-derived strains have more opportunities to recombine relative to strains sampled from geographically and taxonomically separated hosts.

## Supplemental Information

**Figure S1. Recombination among *Bacteroides uniformis* and *Bacteroides humanifaecis* genomes.** Ratio of homoplastic (*h*) to non-homoplastic (*m*) polymorphisms for randomly sampled subsets of *Bacteroides uniformis* genomes and the two available *B. humanifaecis* genomes. The red lines indicate *h/m* values expected when homoplasies arise exclusively from convergent mutations.

**Figure S2.** Ratio of homoplastic to non-homoplastic polymorphisms (*h/m*) for randomly sampled subsets of *Bacteroides fragilis* strains derived from human hosts (Groups 1-10) and non-human hosts (Group ‘nonhuman’). The group labeled ‘nonhuman’ consists of strains derived from domestic pigs (*n* = 3), chickens (*n* = 2), and orangutans (*n* = 10). That *h/m* values remain low for the group of genomes from ‘non-human’ hosts indicates a decreased rate of recombination among these genomes.

**Table S1.** Accession numbers and NCBI species designations for all *Bacteroides* genomes analyzed in this study, along with their inclusion status dependent on whether they were ≥99% complete and contained ≤1% contamination.

**Tables S2–S13.** Accession numbers, GTDB classification, and isolation sources of *Bacteroides* strains with recombination patterns that warranted reclassification.

**Table S14.** Comparison of *h/m* values (present study) and Tmrca /Tmosaic values (12) for seven species of *Bacteroides*.

